# Disentangling genetic effects on transcriptional and post-transcriptional gene regulation through integrating exon and intron expression QTLs

**DOI:** 10.1101/2023.04.27.538308

**Authors:** Anneke Brümmer, Sven Bergmann

## Abstract

Expression quantitative trait loci (eQTL) studies typically consider exon expression of genes and discard intronic RNA sequencing reads despite their information on RNA metabolism. Here, we quantified genetic effects on exon and intron levels of genes and their ratio in lymphoblastoid cell lines, revealing thousands of cis-QTLs of each type. Genetic effects were often shared between cis-QTL types, but 6084 (41%) were not detectable at exon levels. We show that exon levels preferentially capture genetic effects on transcriptional regulation, while exon-intron-ratios better detect those on co- and post-transcriptional processes. Considering all cis-QTL types substantially increased the number of colocalizing GWAS variants (by 61%). It further allowed dissecting the potential gene regulatory processes underlying GWAS associations, suggesting comparable contributions by transcriptional (48%) and co- and post-transcriptional regulation (42%) to complex traits. Overall, integrating intronic RNA sequencing reads in eQTL studies expands our understanding of genetic effects on gene regulatory processes.

## Introduction

Expression quantitative trait loci (eQTLs) are genetic variants associated with gene expression levels. While cis-eQTLs directly affect the expression of nearby genes, trans-eQTLs indirectly modulate the expression of distal genes through affecting nearby regulatory genes or elements. Mapping eQTLs has emerged as a powerful tool to identify functional genetic variants that affect gene expression and has been applied in different cell types, tissues, human populations, during ageing, upon infection, and between sexes ^1–6^. eQTLs were found to colocalise with genetic variants associated with human traits through genome-wide association studies (GWAS), suggesting a causal role for gene expression in mediating such traits ^7,8^. However, the extent of colocalization between GWAS variants and eQTLs was rather small (∼21% of GWAS variants on average per trait ^9^). The reasons could be that sample sizes of eQTL studies are generally smaller than those of GWAS studies, which may not allow resolving eQTLs with weaker effects, that eQTLs may not have been determined in the trait-relevant cell types, or that eQTLs may not capture the trait-relevant gene regulatory processes.

Even though eQTLs provide a strong indication for a genetic variant impacting the regulation of gene expression, the specific regulatory process affected remains ambiguous. It could range from transcription regulation, RNA splicing and processing to the regulation of RNA stability. To overcome this ambiguity, QTLs have been mapped for a variety of molecular phenotypes, such as DNA accessibility, DNA methylation, histone modifications, transcription factor binding, splicing ratios, polyadenylation site usage, ribosome-binding, and protein levels, revealing valuable insights into the genetic effects on specific gene regulatory processes ^1,10–14^. While some of these molecular phenotypes can be quantified from RNA-Seq data, like gene expression for eQTLs, others require data from more advanced experimental high-throughput methods. As a consequence, such QTLs have only been studied for few cell types or tissues, and with relatively small sample sizes, limiting the statistical power for detecting QTLs with smaller effects. Thus, a way to infer the gene regulatory process affected by a genetic variant directly from gene expression measurements would be very valuable for advancing the understanding of the molecular mechanisms of eQTLs.

Over the past decade, it has become clear that intronic RNA-Seq reads contain valuable information about gene regulation. Gaidatzis et al. ^15^ showed that intron expression levels of genes are, like exon expression levels, primarily determined by transcription, while the ratios of exon to intron expression levels, cancelling out transcriptional influences, are more sensitive to post-transcriptional regulation. This approach has been widely applied by others to distinguish between transcriptional and post-transcriptional gene expression changes ^16–18^. Another study demonstrating the value of intronic reads was by La Manno et al. ^19^, who used intronic and exonic RNA-Seq reads to estimate precursor and mature mRNA levels in single cells, allowing them to predict the future transcriptional regimes of individual cells from these, under the assumption that the transition from precursor to mature mRNA (i.e. RNA processing) is constant. This method (termed RNA velocity) has become a standard in single-cell RNA-Seq analyses, and modifications of it have already been proposed ^20–22^. Together, analysing intronic RNA-Seq reads – which are contained in RNA-Seq data, even from polyA-selected RNA ^15^ – on top of exonic reads provides a deeper understanding of gene regulatory processes.

Given these previous breakthroughs, we investigated here the use of intronic RNA-Seq reads for improving the understanding of genetic effects on gene regulation. We determined QTLs for exon and intron expression levels of genes, as well as their ratio using data from 901 lymphoblastoid cell lines (LCLs) from European individuals. We found thousands of genetic variants associated with each of the three gene expression measures, including 41% that were not detectable based on exon expression levels. Considering all QTL types increased the fraction of GWAS variants colocalizing with QTLs (from 19% on average per trait for exon level QTLs to 27% for all three QTL types). Furthermore, we show that integrating the information from all QTL types improves our understanding of the impact of genetic variants on gene regulatory processes and complex traits.

## Results

### Detection of QTLs for exon and intron expression levels and their ratio

To better understand the effects of genetic variants on different gene regulatory processes (Figure 1A), we analysed QTLs for exon expression levels (referred to as exQTLs), and intron expression levels (inQTLs), and their log2 ratio (or, equivalently, their log2-difference; referred to as ex-inQTLs). Exon and intron expression levels for each gene were quantified from RNA-Seq data from lymphoblastoid cell lines (LCLs). We verified that genetic associations with different gene expression measures (referred to as different QTL types below) were reproducible in two independent data sets, the CoLaus ^23^ and European samples of the Geuvadis data set ^2^, despite different sequencing depths and proportions of intronic RNA-Seq reads (see Supplementary Text, Figure S1). To maximise the power for detecting QTLs, we combined the two data sets, resulting in a total of 901 samples. Using this combined data set, we detected significant exQTLs, inQTLs, and ex-inQTLs (FDR < 5%) for 77%, 73%, and 63% of tested genes, respectively, corresponding to 8649, 7590, and 5706 genes (Figure 1B; Supplementary Table S1). We found that the fraction of QTLs located upstream of associated genes, where transcription regulatory regions are preferentially located, was significantly larger among exQTLs (32%) than among the other two QTL types (Figure 1C). In contrast, the fraction of QTLs located within the transcribed gene regions, potentially affecting post-transcriptional regulation, was significantly larger for ex-inQTLs (57%).

**Figure 1:**
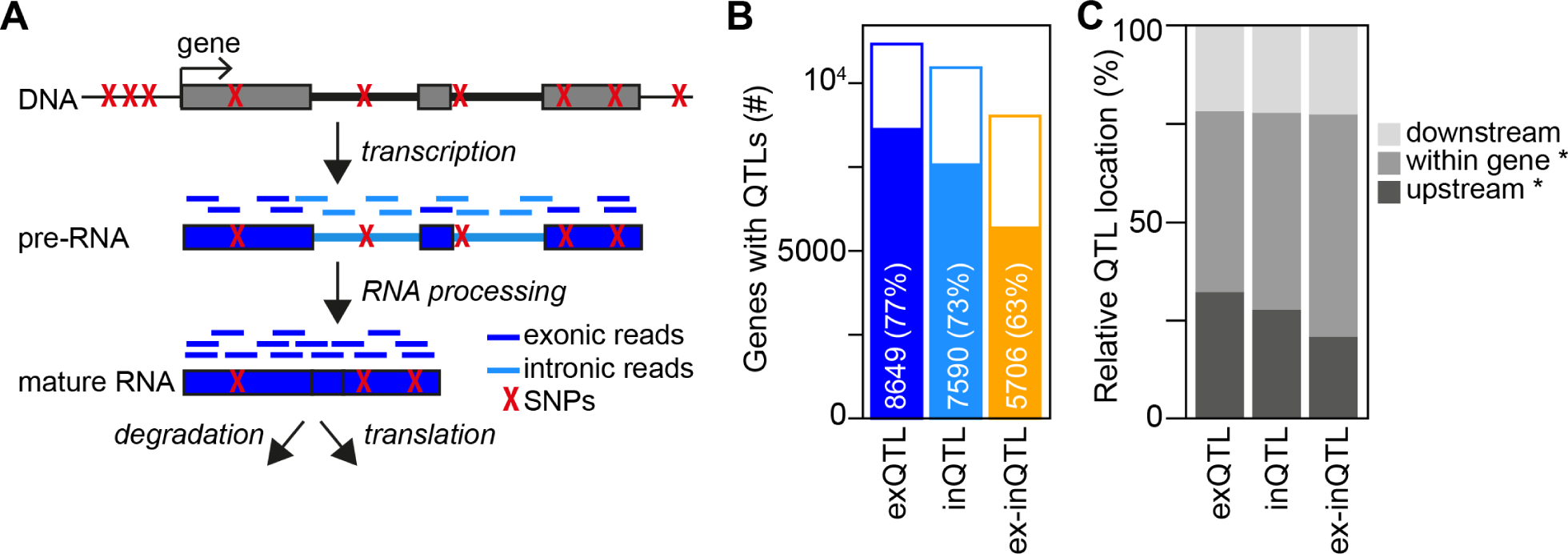
Identification of cis-QTLs for exon and intron expression levels and their ratio. (A) Schematic representation of the steps of gene expression regulation, from DNA over pre-RNA to mature RNA. Genetic variants that could potentially affect different steps are shown in red, and RNA-Seq reads mapping to exons and introns are indicated. (B) Number and percentage of genes with cis-QTLs (filled bars) of tested genes (full bars) for different QTL types. (C) Location of top QTLs relative to their associated genes for different cis-QTL types. The fractions of QTLs upstream of and within genes are each significantly different between QTL types (p<1e-6, Fisher exact test).

### Genetic effects are frequently shared between QTL types

Since for many genes we found several QTL types (6833 out of 10721 genes with any QTLs), we further investigated the sharing of genetic effects between QTL types. We considered QTL effects as shared if the positions of the top QTLs, i.e. the most significantly associated variants for a given gene and for each type, were identical or had similarly strong associations with the gene expression measures with consistent effect directions. This approach revealed substantial sharing (for 6069 genes, including 2230 genes with shared QTL types with identical top QTL variants; Supplementary Table S2). Most sharing occurred between inQTLs and ex-inQTLs (referred to as in&ex-inQTLs), observed for 3327 genes, corresponding to 66% of genes with both such QTLs, followed by exQTLs and inQTLs (ex&inQTLs), observed for 3107 genes, corresponding to 55% of genes with both such QTLs (Figure 2A). Shared exQTLs and ex-inQTLs (ex&ex-inQTLs) were slightly rarer, observed for 2178 genes, corresponding to 43% of genes with such QTLs. 1008 genes presented QTL signals shared between all three QTL types (ex&in&ex-inQTLs; 23% of genes with the three QTL types). The direction of the QTL effects was mostly identical for shared ex&inQTLs, supporting a predominant effect on transcriptional regulation for these (Figure S2A). Overall, combining shared QTL signals, we identified 14869 cis-QTL signals, of which 40.9% were derived from inQTLs or ex-inQTLs and were not detected by exQTLs (Figure 2A).

**Figure 2:**
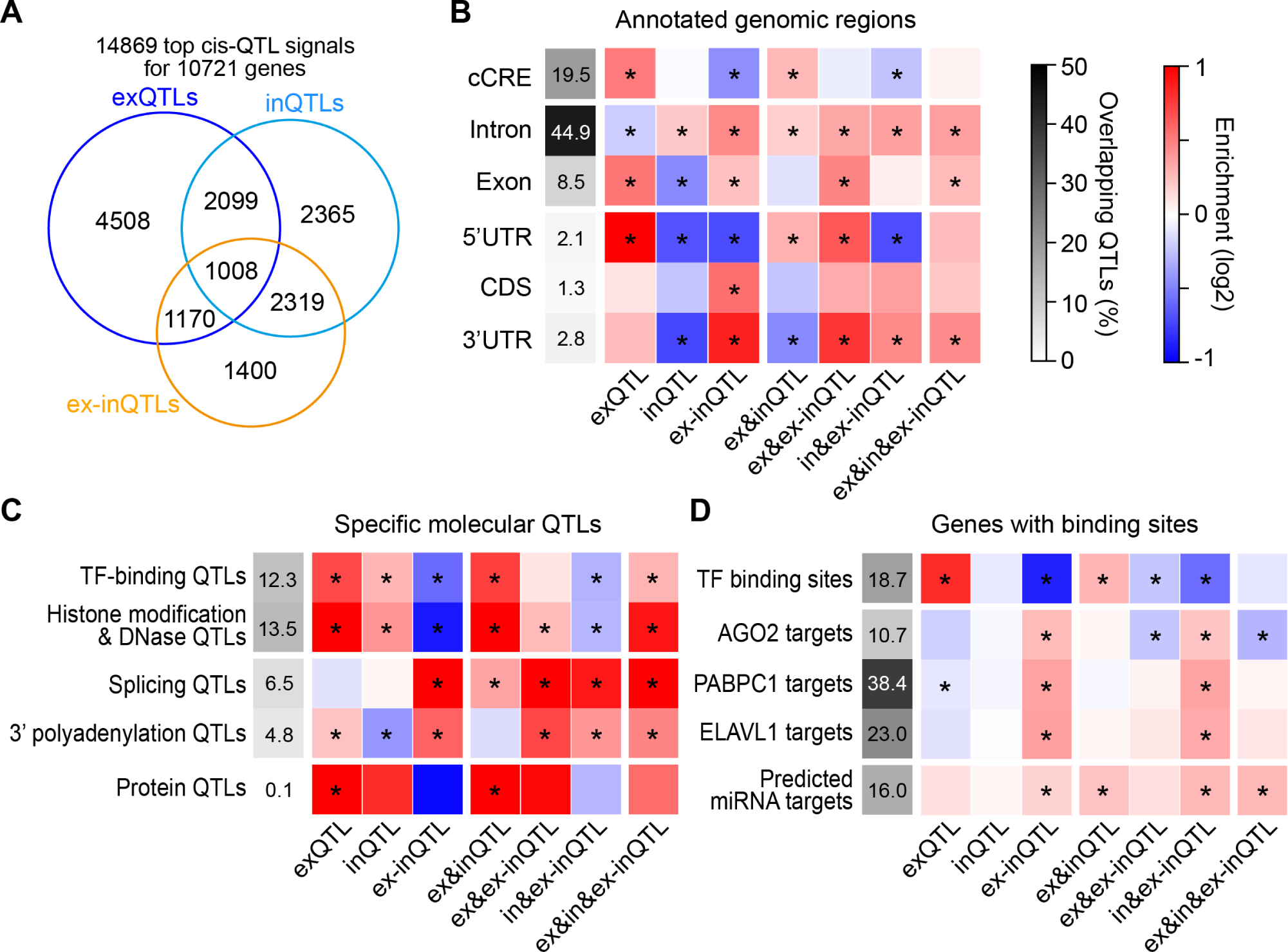
Sharing between top cis-QTL signals, and location within specific genomic regions, overlap with specific molecular QTLs, and with genes with certain binding sites. (A) Venn diagram showing the sharing between different types of cis-QTLs. In total, 14869 independent top cis-QTL signals for 10721 genes were detected. (B) Enrichment within annotated genomic regions (represented as a blue-red heatmap) for different types of cis-QTLs and shared cis-QTLs relative to cis-QTLs not contained in or shared with that group of cis-QTLs. Asterisks indicate a significant enrichment (p<0.05, Fisher exact test). The overall overlap for all cis-QTLs with different genomic regions is indicated in grayscale. cCRE: candidate cis-regulatory element, UTR: untranslated region, CDS: coding sequence. (C) Similar as (B), but for enrichment in overlap with specific molecular QTLs identified for LCLs. (D) Similar as (B), but for enrichment within genes harboring certain binding sites determined in LCLs, considering top cis-QTLs located within genes.

In our study, effects shared between QTL types beyond top QTLs – that is, between conditionally independent QTLs detectable after regressing out any stronger masking QTL signals – was relatively rare. Indeed the majority of genes (76%, 71%, and 66% for exQTLs, inQTLs, and ex-inQTLs, respectively) presented only one (the top) QTL signal (Figure S2B; Supplementary Table S4). Nevertheless, we found a small but significant bias towards top ex-inQTL and inQTL signals being more frequently shared with secondary exQTL signals (6.6% and 6.1%, respectively) than the opposite (top exQTL signals shared with secondary ex-inQTL or inQTL signals: 4.7% and 3.9%; p<0.05, Fisher exact test; Figure S2C). This may indicate that post-transcriptional effects on exon levels (detected by top ex-inQTLs) can be masked by other, stronger regulatory processes acting on exon levels, but these cases are rare.

Interestingly, we also detected shared effects between top cis-QTL signals of the same type but for different genes (Supplementary Table S2). Sharing of top exQTLs (12.8% of all genes with exQTLs) and inQTLs (11% of genes with inQTLs) was significantly more frequent than between ex-inQTLs (6.6% of genes with ex-inQTLs; p < 1e-18, Fisher exact test). This agrees with a prevalent transcriptional co-regulation of neighbouring genes, as already reported ^24^, while post-transcriptional regulation (detected by exon-intron ratios) appears more gene-specific.

### Different QTLs types and shared QTLs are enriched within genomic regions linked to distinct gene regulatory processes

To understand in more detail which gene regulatory processes are affected by different QTL types and shared QTLs, we analysed their location within specific genomic regions and sites.

As a first indication of their regulatory function, we examined the QTL location within annotated genomic regions (Figure 2B). Compared to other QTL types, exQTLs were enriched in transcription regulatory DNA elements, i.e. candidate cis-regulatory elements (cCREs) for B lymphocytes, and in 5’ untranslated regions (UTRs) of the associated genes. In contrast, ex-inQTLs were depleted in these regions and were enriched in 3’UTRs, coding sequences (CDSs) and intronic regions. inQTLs were enriched in intronic regions but depleted in exonic regions. Shared ex&inQTLs showed enrichment in cCREs and 5’ UTRs, but not in 3’UTRs. Shared ex&ex-inQTLs were generally enriched in transcribed gene regions, particularly in 5’ and 3’ UTRs. Shared in&ex-inQTLs were depleted in cCREs and 5’ UTRs and enriched in introns and 3’ UTRs. QTLs shared between all three types were also enriched in transcribed gene regions, with a slight preference for 3’ UTRs and introns.

Next, we analysed the overlap with specific molecular QTLs identified previously for LCLs (Figure 2C; see Methods). We found strong preferential overlaps for exQTLs and shared ex&inQTLs with QTLs for transcription factor (TF) binding, histone modifications and DNA accessibility, and protein levels. In contrast, ex-inQTLs and shared in&ex-inQTLs were depleted for overlaps with such QTLs and instead preferentially overlapped with QTLs for splicing and 3’ polyadenylation. Shared ex&ex-inQTLs and ex&in&ex-inQTLs overlapped also with these QTLs and additionally with histone and chromatin QTLs, suggesting a role of these shared QTLs in one or several of these processes.

Finally, we investigated the enrichment of groups of QTLs within measured TF-binding sites and experimentally identified target genes of post-transcriptionally regulatory RNA-binding proteins (RBPs), AGO2, PABPC1 and ELAVL1, in LCLs, and predicted target genes of microRNAs (miRNAs) highly expressed in LCLs (Figure 2D). exQTLs and shared ex&inQTLs showed strongest enrichment for TF-binding sites, while ex-inQTLs and shared ex&ex-inQTLs and in&ex-inQTLs were depleted in these regions. Instead, ex-inQTL and in&ex-inQTLs were enriched within genes with RBP- and miRNA-binding sites.

In summary, the enrichment analysis of QTL groups within specific genomic regions indicates that QTLs for exon expression levels are enriched for transcriptional effects, while ex-inQTLs rather represent effects on post-transcriptional processes. Shared QTLs seem to further dissect the diverse effects of QTL types on gene regulatory processes.

### TFs have stronger trans-effects on exon levels, while RBPs have stronger trans-effects on exon-intron ratios

Our enrichment analysis focussed on top cis-QTLs, which have a high probability for being causal, but may not always be, due to strong genetic correlation between neighbouring genetic variants leading to similar significant (sometimes indistinguishable) associations with the gene expression measurements. To further investigate how the effects of different gene regulatory processes are captured by the three QTL types in our study, we investigated trans-effects - on genes located on different chromosomes or at distances larger than 5 million base pairs - of regulatory factors (Figure 3A). As regulatory factors we considered 721 TFs and 691 RBPs with any type of cis-QTL association in our data set, and 47 miRNAs with cis-QTLs previously identified in LCLs from European individuals (Lappalainen et al. 2013). We identified trans-QTLs of all types (trans-exQTL, trans-inQTL, trans-ex-inQTL) for cis-QTLs of these regulatory factors, and detected 399 unique trans-QTL associations between a regulatory factor and a potential target gene (183 for TFs, 131 for RBPs and 85 for miRNAs; Supplementary Table S5). While TFs had larger proportions of trans-associations with exon and intron levels (36% and 42%, respectively) than with their ratio (22%; Figure 3B), RBPs and miRNAs had higher proportions of trans-associations with intron levels (46% and 54%, respectively) and exon-intron-ratios (34% and 28%, respectively) than with exon levels (20% and 18%, respectively). Trans-exQTL associations were significantly more frequent for TFs than for RBPs and miRNAs, while trans-ex-inQTL associations were significantly more frequent for RBPs than for TFs. Notably, the proportions of different types of trans-QTL associations for TFs and RBPs were similar when considering only cis-QTLs of any one type (Figure S3A), indicating that all types of cis-QTLs of regulatory factors enable similar functional trans-effects.

**Figure 3:**
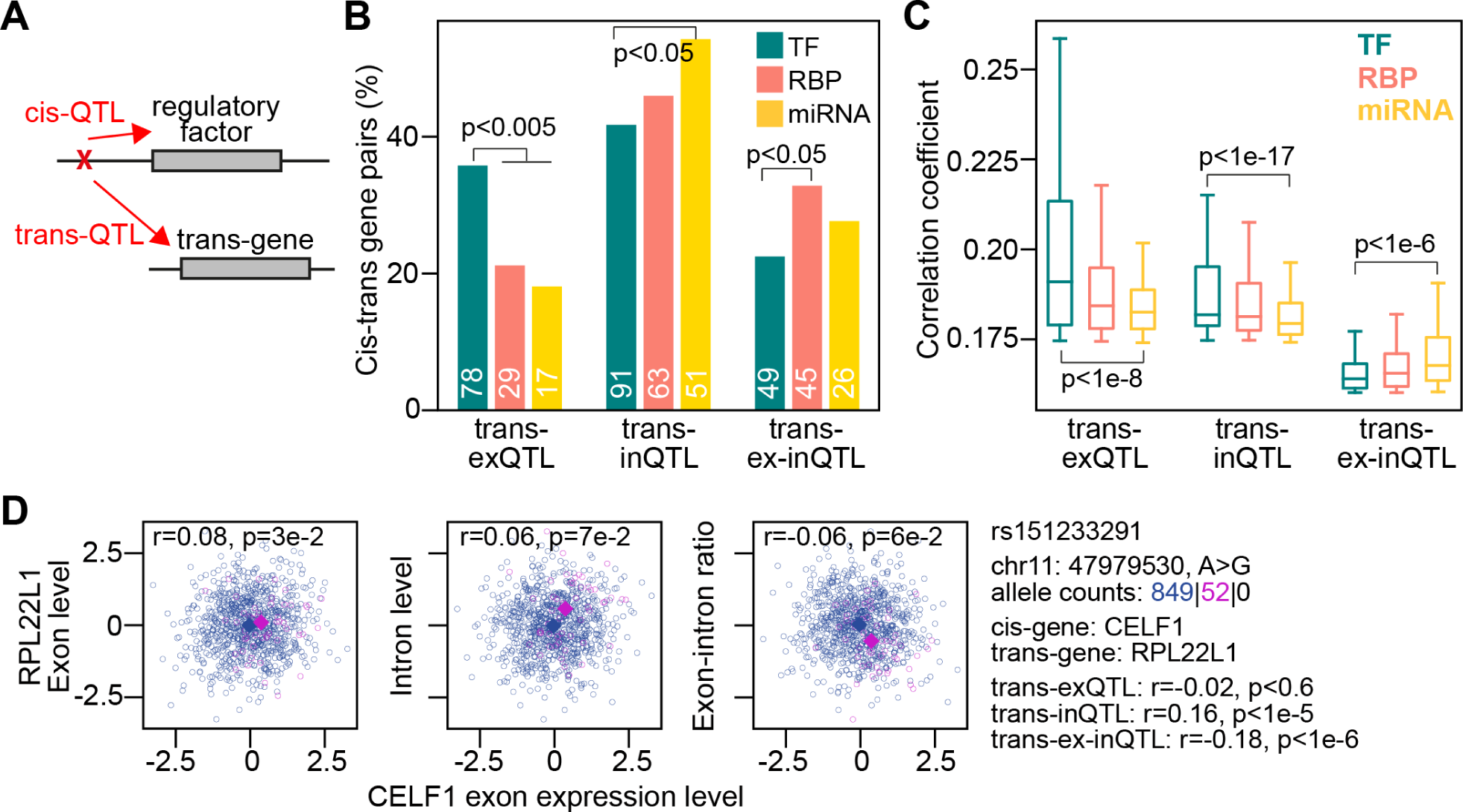
Trans-QTL effects on exon or intron levels or their ratio for cis-QTLs of regulatory factors. (A) Illustration of a cis-QTL for a regulatory factor that also has a trans-QTL effect on a (target) gene located at a distance of at least 5Mb or on a different chromosome, which indicates a potential causal relationship between the cis-associated regulatory factor and the trans-associated gene. (B) Percentages of detected trans-QTL associations of different types for transcription factors (TF), RNA-binding proteins (RBP), and miRNAs. The numbers of trans-QTL associations are indicated at the bottom of each bar. P values are calculated using Fisher’s exact test. (C) Boxplots of absolute correlation coefficients (comparable to effect sizes) for significant trans-QTL associations of different types shown in (B). P values are calculated using a Ranksum test. (D) Example for a cis-regulated RBP, CELF1, associated in trans with RPL22L1. Shown are scatter plots of the RBP’s normalised exon levels (x-axis) and the normalised exon levels (left panels), intron levels (middle panels) and their ratio (right panels) of the trans-associated gene (y-axis). Further information on the QTL variant and trans association are indicated on the right. Correlation coefficients between the expression levels/ratios of the RBPs and their trans-associated genes are indicated inside each panel. Circles of different colors represent individuals with different genotypes, and colored squares indicate the median values for individuals with that genotype.

Comparing the strengths of significant correlations between genotype and gene expression measurements across regulatory factors, we further found, that cis-QTLs of TFs were generally stronger correlated with exon and intron levels of trans-genes than cis-QTLs of RBPs or miRNAs. In contrast, the correlation with exon-intron-ratios of trans-genes was stronger for cis-QTLs of RBPs and miRNAs than for those of TFs (Figure 3C).

Examples for trans-QTL associations with cis-regulated RBPs are between DGCR8 (involved in processing of primary miRNA transcripts) and exon or intron levels of miRNA harbouring genes, between LSM11 (involved in histone mRNA 3’-end processing) and exon or intron levels of histone genes, and between between CELF1 (implicated in splicing and stability regulation of target RNAs) and intron levels and exon-intron ratios of RPL22L1 (Figures 3D and S3B,C). Notably, genetic trans-associations were mostly not detectable as significant correlations between the gene expression measurements of cis- and trans-regulated genes. The miRNAs with most trans-associations were miR-1255a and miR-550a (8 associations each), which had 9 trans-associations with intron levels, 7 with exon-intron-ratios, and 3 with exon levels of associated genes. Of these 6, 5, and 0 genes had predicted miRNA target sites in their 3’ UTR, respectively ^25^.

In summary, the trans-association analysis confirms that effects on post-transcriptional regulatory processes are better detectable at exon-intron-ratios, while transcriptional regulation is better detectable through exon levels. Intron levels appeared sensitive to both types of regulation.

### inQTLs and ex-inQTLs substantially increase the colocalization with GWAS variants

Having developed a good understanding of the regulatory processes affected by different QTL types and shared QTLs, we next investigated if combining the information from the three QTL types might aid the understanding of the functional impact of GWAS variants.

We first identified QTLs that colocalized with GWAS variants. We used the regulatory trait concordance (RTC) method ^8^, which tests for co-localization between variants within the same genomic regions surrounded by recombination hotspots (see Methods). Of the tested top cis-QTLs, the percentage that colocalized with GWAS variants was similar for different QTL types (30-34%), supporting their similar functional relevance for human traits (Figure 4A). Adding inQTLs and ex-inQTLs increased the number of GWAS variants colocalizing with QTLs from 3269 for exQTLs (18.3% of tested GWAS variants) to 5276 (25.1% of tested GWAS variants) for all QTL types (Figure 4B). Thus, the number of colocalizing GWAS variants increased by 61%. For 89 GWAS traits with at least 20 colocalizing variants, colocalization increased to 26.6% of tested GWAS variants on average per trait (Figures 4C and S4A) from 19.5% for colocalization with exQTLs only. Among GWAS traits with a large amount of colocalization were skin-related traits (tan response, basal cell carcinoma) and lung function traits, while traits with a low colocalization included cholesterol, apolipoprotein, alkaline phosphatase, and non-albumin protein traits. GWAS traits with a relatively large colocalization with exQTLs included testosterone, white blood cell traits (eosinophil and monocyte percentage), cognitive and neurological traits (cognitive ability, intelligence, schizophrenia), and body fat, while GWAS traits with relatively large colocalization with inQTLs or ex-inQTLs (and low colocalization with exQTLs) included lung function traits, heart-related traits (PR interval, atrial fibrillation) and a liver-related trait (gamma-glutamyl transferase). Many GWAS variants (1952 or 37%) colocalized with several QTL types (Figure S4A), potentially providing valuable information on the type of regulatory process contributing to a complex trait. Overall, including inQTLs and ex-inQTLs increased the number of colocalizations with GWAS traits substantially.

**Figure 4:**
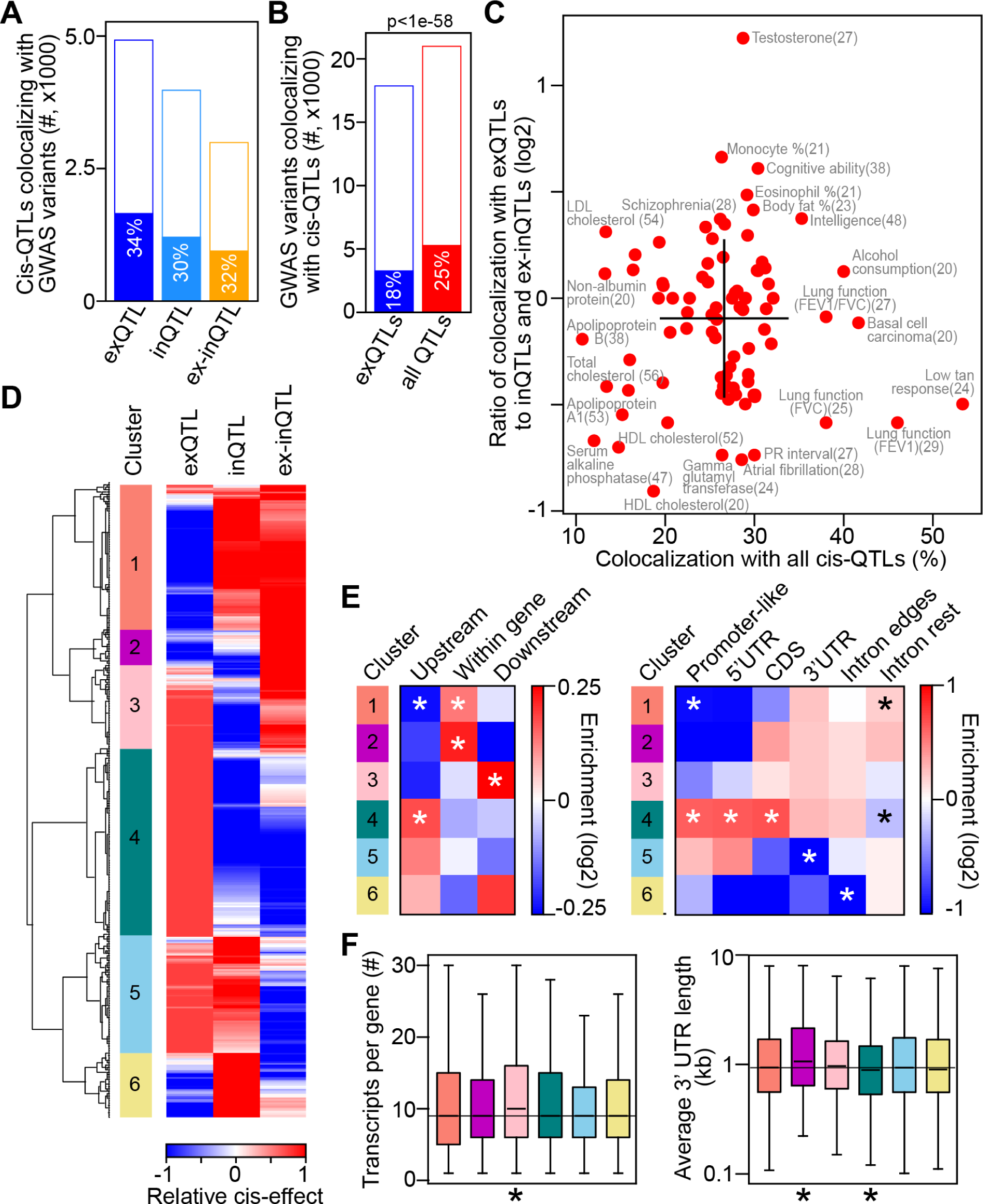
Colocalization between GWAS variants and top cis-QTLs of different types. (A) Number of top cis-QTLs colocalizing with GWAS variants (filled bars) of tested QTLs (full bars) for different QTL types. (B) Number of GWAS variants colocalizing with top cis-QTLs (filled bars) of tested GWAS variants (full bars) for colocalization with exQTL and all QTL types. P value was calculated using Fisher’s exact test. (C) Percentage of colocalizing GWAS variants (x-axis) and log2 ratio of GWAS variants colocalizing with exQTL to those colocalizing with inQTLs or ex-inQTLs (y-axis), for 89 GWAS traits with at least 20 colocalizations. GWAS traits with extreme values are labeled, and the number of colocalizing GWAS variants is indicated in parenthesis. Black lines indicate the mean +/- standard deviation for each axis. (D) Clustering of normalized cis-QTL effects for 2486 QTLs colocalizing with GWAS trait variants and with measured effects for all types of QTL-associations. Six distinct clusters are labeled. (E) Enrichment of QTLs upstream, within or downstream of associated genes (left panel) and enrichment within annotated genomic regions (right panel) for cis-QTLs of each cluster compared to cis-QTLs of other clusters. Intron-edges include 100 bps at intron starts and ends. *, p<0.05, calculated using Fisher exact test. (F) Number of annotated transcripts (left panel) and average 3’ UTR length (right panel) of genes associated with top cis-QTLs in different clusters (indicated by color code from (D)). *, p<0.05, calculated using Ranksum test compared to cis-QTLs in other clusters.

### Dissecting the gene regulatory processes underlying GWAS associations

Next, we examined if the quantified cis-effects on exon and intron levels and exon-intron ratios of QTLs colocalizing with GWAS variants might help elucidate the gene regulatory mechanisms underlying GWAS associations. We focussed on 2486 top cis-QTLs that colocalized with GWAS variants and for which all three types of cis-effects on the target gene’s gene expression measures were quantified, including at least one significant cis-QTL association. After normalising absolute cis-effects between QTL types and across variants, we performed hierarchical clustering with these “relative” cis-effects and obtained six clusters with distinct patterns of cis-effects (Figure 4D). Most variants were in cluster 4, strongly affecting exon levels (29.5%), followed by cluster 1 (22.9%), affecting intron levels and exon-intron ratios, and cluster 5 (18.6%), affecting exon and intron levels but not their ratios. Some GWAS traits colocalized with QTLs that were enriched for QTLs from a certain cluster, e.g. lung function-related traits for QTLs in cluster 3 (affecting exon levels and exon-intron ratios) or eosinophil-related traits for QTLs in cluster 4 (affecting exon levels; Figure S4B).

The results presented before suggest that variants in clusters 4 and 5 likely affect transcriptional processes, while variants in clusters 1, 2 and 3, with comparably strong effects on exon-intron-ratios, likely affect splicing and other post-transcriptional processes. To examine this hypothesis, we analysed the locations of the QTLs in each cluster and the structural properties of their associated genes. Indeed, variants in cluster 4 were enriched in promoter-like elements and generally upstream and at the beginnings of genes (Figure 4F). Variants in cluster 5 were also preferentially located upstream and at the beginnings of genes and, additionally, were depleted in 3’ UTRs. In contrast, variants in clusters 1 to 3 were depleted in promoters, upstream of genes, and at the beginning of genes (5’UTRs), but were, in general, enriched within the transcribed gene regions and downstream of genes (cluster 3). Finally, we examined the structural properties of the genes regulated by cis-QTLs in different clusters. Variants in cluster 3 were associated with genes with significantly more annotated transcripts, potentially indicating a more complex RNA processing for these genes, compared to genes associated with QTLs belonging to other clusters. Genes regulated by variants in cluster 2 had significantly longer 3’ UTRs, potentially indicating that these genes are under more extensive post-transcriptional regulation, while genes regulated by variants in cluster 4 had significantly shorter 3’ UTRs. Thus, the analysis of QTL location and associated genes’ structural properties supports the hypothesis that variants in clusters 4 and 5 likely affect transcriptional processes, while variants in clusters 1 to 3 likely affect splicing and post-transcriptional processes.

Altogether, combining the information from different QTL types improves the understanding of the regulatory processes underlying GWAS associations.

### Examples for genetic effects on post-transcriptional and transcriptional gene regulation with relevance for complex traits

An example for a cis-QTL likely affecting co- or post-transcriptional gene regulation and colocalizing with a GWAS trait is rs60252802, a cis-QTL from cluster 1 (Figure 5A). This cis-QTL is associated with intron level and exon-intron-ratio of SDF4, a calcium-binding protein involved in regulating calcium-dependent cellular activities ^26^. It is located in the first intron of SDF4 and colocalizes with GWAS variants for systemic lupus erythematosus, an autoimmune disease affecting multiple organs. Inspecting the RNA-Seq read distributions of homozygous individuals with reference and alternative genotypes indicates that this cis-QTL appears to increase the probability for mis-splicing, through introducing an alternative 5’ splice site, leading to an extension of the first exon (which is part of the 5’ UTR) by almost 400 nucleotides. Another example is rs8031627, a cis-QTL from cluster 2 (Figure 5B). It is associated with the exon-intron-ratio of SMAD3, a transcription factor functioning in the transforming growth factor-beta signalling pathway, and colocalizes with GWAS variants for the lung function trait FVC. The top cis-QTLs for exon and intron levels do not colocalize with the GWAS trait variants. The top cis-ex-inQTL is located in the last exon, which contains the ∼5000 nucleotides long 3’UTR. Although the top cis-ex-inQTL is not directly located within a predicted miRNA binding site, it is still possible that this QTL, or seven other cis-ex-inQTLs sharing the QTL effect and also located in the 3’UTR, interfere with miRNA targeting in their vicinity, or with binding of other post-transcriptional regulatory factors to the 3’UTR. An example for a cis-QTL from cluster 3 is rs2711977 (Figure 5C), which is associated with exon levels and exon-intron-ratios of TMEM156, a transmembrane protein, and colocalizes with GWAS variants for monocyte count. TMEM156 expression is also affected by an inQTL, which is not shared with the two other QTL types and did not colocalize with variants of this GWAS trait. While the colocalizing top cis-QTL is upstream of the gene, another QTL (rs2254075), sharing the exQTL and ex-inQTL signals with the top cis-QTL, is located in the second exon, a 150 nucleotide long exon that is contained only in non-coding transcript annotations of TMEM156. Only individuals homozygous for the minor allele express this exon, and they also exhibit lower expression of all other exons of that gene compared to individuals homozygous for the major allele. It is possible that inclusion of this exon, which contains stop codons in all reading frames, triggers nonsense-mediated decay of that transcript, leading to its reduced overall expression. Alternatively, the two QTLs (upstream and in the second exon) together lead to simultaneous changes in splicing of that exon and the overall transcription rate.

**Figure 5:**
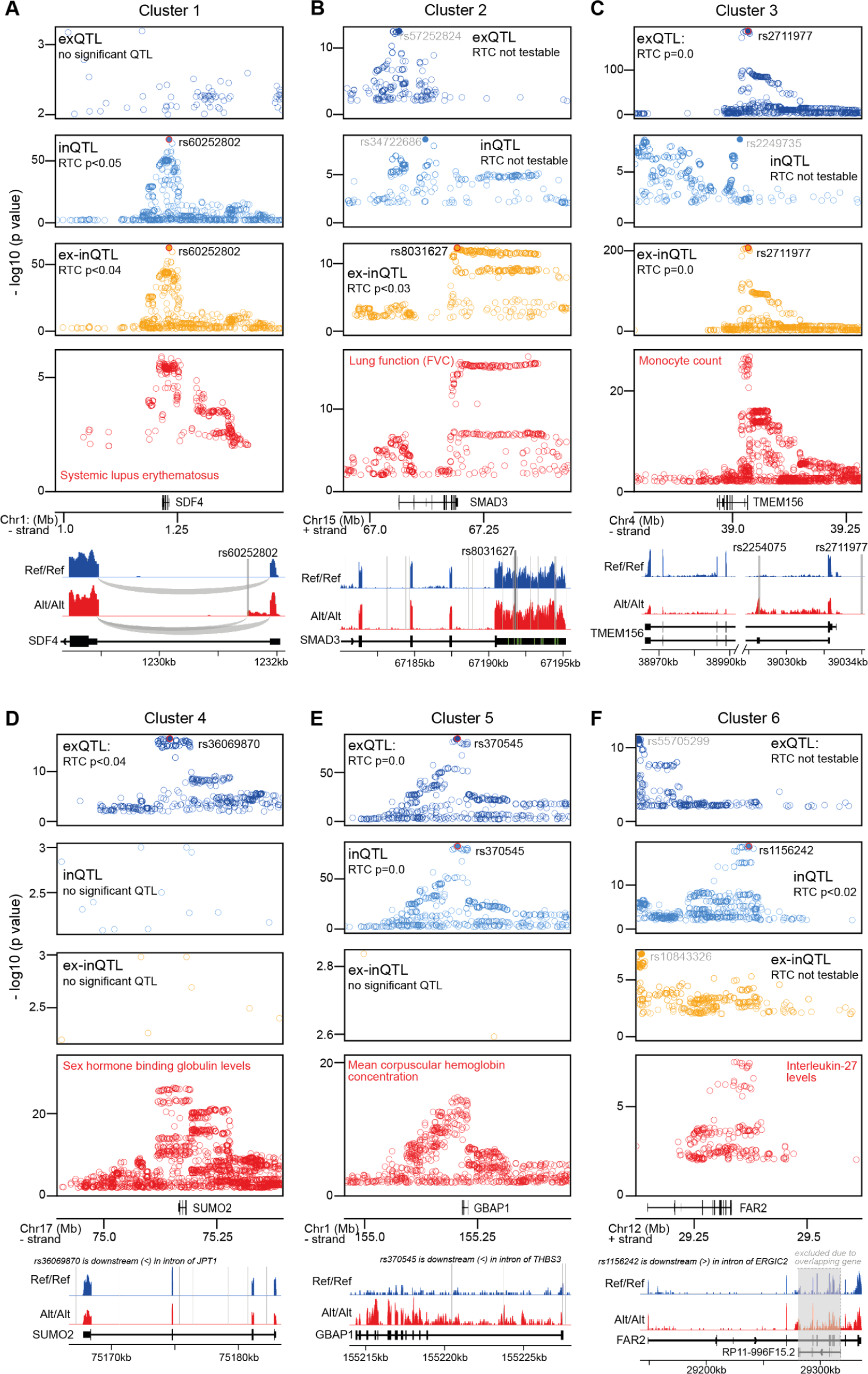
Examples for cis-QTLs from different clusters colocalizing with GWAS variants. In each subfigure, the top three panels show the -log10 nominal p-values (<0.01), in a region of 500kb around the top cis-QTL (x-axis), for cis-QTL associations with exon levels (dark blue), intron levels (blue) and exon-intron-ratios (orange), and the fourth panel shows the -log10 p-values for the GWAS trait associations (red) in the same region. The rsID of the top cis-QTL(s) are indicated (in black, if colocalized with the GWAS trait variants, or in gray if not). Colocalization via the RTC method (see Methods) was testable for QTLs and GWAS variants within the same genomic region between recombination hotspots. The bottom panel shows examples for RNA-Seq read distributions at the associated gene from two homozygous individuals, one with reference (Ref/Ref; blue) and one with alternative (Alt/Alt; red) genotype, for the cis-QTL variant. The positions of the top cis-QTL as well as cis-QTLs sharing the QTL signal are indicated with thick or thin lines, respectively. (A) A cis-QTL from cluster 1, associated with SDF4 and colocalizing with a GWAS variant for systemic lupus erythematosus. (B) A cis-QTL from cluster 2, associated with SMAD3 and colocalizing with a GWAS variant for FVC, a trait related to lung function. (C) A cis-QTL from cluster 3, associated with TMEM156 and colocalizing with a GWAS variant for monocyte count. (D) A cis-QTL from cluster 4, associated with SUMO2 and colocalizing with a GWAS variant for sex hormone binding globulin levels. (E) A cis-QTL from cluster 5, associated with GBAP1 and colocalizing with a GWAS variant for mean corpuscular hemoglobin concentration. (F) A cis-QTL from cluster 6, associated with FAR2 and colocalizing with a GWAS variant for interleukin-27 levels.

Examples for cis-QTLs likely modulating transcriptional gene regulation and mediating a complex trait are rs36069870 from cluster 4, associated with exon levels of SUMO2, a small ubiquitin-like protein modifier, and colocalizing with GWAS variants for sex hormone binding globulin levels, and rs370545 from cluster 5, associated with exon and intron levels of GBAP1, a pseudogene potentially regulating the expression of its related coding gene beta-glucosylceramidase 1 through sponging of miRNAs ^27^, and colocalizing with GWAS variants for mean corpuscular hemoglobin concentration. Both of these top cis-QTLs are located in introns of downstream genes on the same strand, JPT1 and THBS3, respectively. A difference between these two examples is that intron levels remain unaffected for SUMO2 (cluster 4), while they are similarly affected as exon levels for GBAP1 (cluster 5). This could indicate that splicing is not rate-limiting for mature RNA levels for cluster4, while the splicing rate limits the transcriptional regulation of mature RNA levels for cis-QTLs in cluster 5. Finally, an example for a cis-QTL from cluster 6 is rs1156242, associated with intron levels of FAR2, encoding a fatty acid reductase enzyme, and colocalizing with a GWAS variant for interleukin-27 levels. It is located in an intron of a downstream gene on the opposite strand, ERGIC2, whose exon and intron levels are both also affected by this cis-QTL. This indicates a likely co-transcriptional regulation of these neighbouring genes.

Further examples for QTLs from each cluster are shown in Figure S5.

## Discussion

In this study we showed that combining traditional eQTL analysis based on exon levels with the analysis of QTLs for intron levels and exon-intron-ratios increases the number of genetic variants associating with gene expression measurements and expands the understanding of their effects on gene expression regulatory processes.

We found that the most significant exQTLs (which are identical to the traditionally analysed eQTLs) often affect transcriptional gene regulation, which, being a pivotal step in gene regulation, has accordingly large effects on gene expression. Smaller gene expression changes by non-transcriptional processes are less detectable by traditional eQTLs. To more comprehensively study the impact of genetic variants on gene expression, the analysis of conditionally independent eQTLs has been proposed ^1,28^, which are detected after stepwise regressing out from the exon expression levels stronger, masking eQTL effects. Conditional eQTLs are expected to have the ability to capture multiple independent genetic effects on gene expression, potentially including subtle effects on gene expression. Our analysis showed that conditional exQTLs do not generally correspond to the gene regulatory effects detected by inQTLs or ex-inQTLs (Figure S2), and thus cannot replace them. This indicates that taking into account intronic RNA-Seq reads provides information on gene expression regulation that is not accessible based on exonic reads only.

Previously, the colocalization of GWAS variants with eQTLs was found to be limited (around 21% per trait for eQTLs from 54 tissues) ^9^. In our data set, the colocalization of tested GWAS variants with traditional eQTLs from LCLs was 19.5% per trait. Adding inQTLs and ex-inQTLs increased the colocalization to 26.6% of tested GWAS variants, thus by 36.4%. Integrative analysis of the effects of all QTL types suggested that 48.1% of colocalizing QTLs likely function in transcription regulation (QTLs in clusters 4 and 5 of Figure 4D), while a similarly large fraction of QTLs (41.8%) had strong effects on exon-intron ratios (clusters 1, 2 and 3), indicating a likely function in post-transcriptional regulation. Thus, the contributions of transcriptional and post-transcriptional processes to complex traits appear to be similar, indicating that both types of regulation are equally functionally relevant, despite smaller effects by post-transcriptional regulation on exon levels. Indeed, QTLs with small effects on exon expression levels of genes can have important functional consequences, such as through promoting exon skipping or inclusion, or alternative splice site usage (Figures 5 and S5) ^29^.

In agreement with previous results on changes in exon and intron expression levels between conditions, and the difference between these changes ^15^, we found that the most complementary effects were between those of exQTLs and ex-inQTLs, as indicated by their enrichment in transcription regulatory sites and regions with post-transcriptional regulatory sites (e.g. 3’ UTRs or genes with target sites of RBPs and miRNAs), respectively. Compared to these two QTL types, inQTLs did not show strong preferences for either of these regulatory regions and similarly, trans-effects on intron levels appeared comparable between TFs and RBPs. While we expect that transcription affects intron levels, less is known about their regulation after transcription or potential regulatory roles of introns. Our analysis suggests that introns are similarly sensitive to regulation by RBPs and miRNAs, as indicated by frequent trans-effects of these on intron levels. Assuming that these effects are direct, this would suggest that post-transcriptional regulators modulate some spliced introns in the nucleus or in the cytoplasm, where a functional role of introns is unclear. To advance the understanding of potential functional roles of some introns, analysing QTLs for levels of single introns might be informative.

The sharing of top cis-QTLs indicated a sizable number of QTLs with shared effects on all three gene expression measures (Figure 2). These QTLs had enriched overlaps with multiple specific molecular QTLs, in particular those modulating transcription regulatory processes and those affecting RNA processing, potentially indicating co-regulated effects on transcription and RNA splicing or polyadenylation, which has been described before ^30^.

Our approach to incorporate intronic reads identified thousands of genetic variants with effects on gene regulation. Further methodological improvements may be possible to enhance the sensitivity of detecting QTLs, such as explicitly modelling the unequal effects of polyA-selection of the sequenced RNA on exon and intron levels, or refining the gene regulatory processes affected by taking into account the QTL effect directions. To allow other researchers to use and further investigate the QTLs identified in this study with LCLs from 901 individuals of European descent, we provide them as Supplementary Material.

We suggest the presented approach to be routinely integrated into eQTL analyses, as it is easy to implement, uses existing RNA-Seq data, and enables an expanded view of genetic effects on gene regulation.

## Material and Methods

### Quantification of gene expression in the CoLaus and Geuvadis data sets

The CoLaus data set ^23^ was available in-house, while the Geuvadis data set ^2^ was downloaded from ArrayExpress (https://www.ebi.ac.uk/arrayexpress/experiments/E-GEUV-3/). We aligned paired-end unstranded RNA-Seq reads from both data sets to the human genome (build GRCh38) using STAR 2.7 ^31^ with transcript annotations from GENCODE ^32^ (version 34 downloaded from www.gencodegenes.org). We counted separately RNA-Seq reads that mapped uniquely to annotated exons and introns of genes using htseq-count ^33^. We considered as intronic all genic regions that were not annotated as exons in any of the gene’s annotated transcripts. We shrank intron annotations additionally by 10 nucleotides at exon-intron boundaries to account for inaccuracies in splice site annotations, as done by Gaidatzis et al. ^15^. As the RNA-Seq data were unstranded, we further created custom gene annotation files which contained only exonic or intronic parts of genes that did not overlap with any other annotated gene using bedtools ^34^. Our exon and intron gene annotation files are available as Supplementary Material. We counted as exonic the reads fully contained within the custom exon annotations (-m intersection-strict) to enrich for reads from mature RNAs. We considered as intronic all reads that overlapped our custom intron annotations (-m union), reasoning that any overlap with an intron indicates its presence. For QTL associations, we only tested genes with a median number of >10 mapped reads across individuals, whether for exonic reads (in case of exQTLs), intronic reads (in case of inQTLs) or for both (in case of ex-inQTLs). We quantified gene expression as RPKM (reads per kilobase per million reads) separately for exonic and intronic reads, using gene lengths calculated from the respective modified gene annotation files. We added to the RPKM levels a pseudo count of 0.1 and log2-transformed them. We calculated exon-intron ratios as the difference between the log2 exon and log2 intron RPKM levels.

### Mapping of cis-QTLs

We restricted our analysis to SNVs with a minor allele frequency >1% and a minor allele count >10. SNV positions were lifted from hg19 over to hg38 using picard-tools LiftoverVcf (https://broadinstitute.github.io/picard/). We restricted our analysis to European individuals or those with European origin, and excluded 89 samples from the Geuvadis data set that were from African individuals (from Yoruba in Nigeria; labelled YRI). For the combined data set, we merged genotypes from the CoLaus and Geuvadis data sets using vcf-merge ^35^.

We used the genotypes’ first three principal components (PCs) as covariates, and the gene expression’s PCs to account for technical and other co-variabilities in the RNA-Seq data, as suggested before ^36,37^. We optimised the number of PCs used as covariates by maximising the number of genes with significant (FDR <5%) cis-QTLs, i.e., the first 70 PCs for exQTLs, the first 50 PCs for inQTLs, and the first 40 PCs for ex-inQTLs. Using that many PCs outperformed using age, sex, or the fraction of intronic reads explicitly as covariates. Notably, the number of PCs that was optimal was the same for all data sets (CoLaus, Geuvadis without YRI, and the combined data set of both). For each gene, we tested all variants located within one million nucleotides (upstream and downstream) of annotated gene starts.

We mapped cis-QTL with QTLtools 1.3 ^37^ to obtain: (1) the top cis-QTL association for each gene after empirically adjusting p values using 1000 permutations, (2) all nominally significant associations, and (3) conditional cis-QTL associations after stepwise regressing out stronger, masking cis-QTL signals from the gene expression measurements.

### Sharing of cis-QTLs

We defined two cis-QTL effects as shared when the top cis-QTL positions were identical or when both top QTLs were within the first conditional QTL signal of the other with consistent effect directions. The first conditional QTL signal is composed of genetic variants that have similar, but less significant, effects on gene expression measures as the top cis-QTL (Supplementary Table S3). We used QTLtools ^37^ to identify the genetic variants belonging to first conditional cis-QTL signals, and to identify conditional QTL signals that are masked by stronger cis-QTL effects. These are detected after stepwise regressing out the effects from stronger QTL signals from the gene expression measurements. We used the same definition of sharing to investigate sharing between conditional cis-QTL signals of ranks 1 to 5 (identified after stepwise regressing out up to four stronger cis-QTLs signals), between top cis-QTLs of the same type but for different genes (to investigate co-regulation of neighbouring genes), and between top cis-QTLs of the same type and for the same gene, but identified in different data sets (to investigate the reproducibility of top cis-QTL signals in the CoLaus and Geuvadis (without YRI) data sets).

### Overlap with annotated genomic regions, specific molecular QTLs and genes with regulatory sites

Genomic regions of exons, introns, coding regions, 5’ UTRs, and 3’ UTRs were taken from GENCODE gene annotations (version 34) ^32^. We downloaded from ENCODE ^38^ annotations of candidate cis-regulatory elements (cCRE) identified through patterns of chromatin modifications and DNA binding factors, and retained only cCREs identified for B lymphocyte cell lines. We considered the following specific molecular QTLs previously identified in LCLs: TF-binding QTLs ^10^, histone modification and DNase hypersensitivity QTLs ^11^, splicing QTLs ^1^, 3’ polyadenylation QTLs ^13^, and protein level QTLs ^12^. QTL positions were lifted over to hg38 using the liftOver tool and liftOver chains from UCSC (http://hgdownload.soe.ucsc.edu). We considered the following regulatory sites experimentally determined in LCLs: TF binding clusters identified using ChIP-Seq (track: TF Clusters downloaded from UCSC Table Browser: https://genome.ucsc.edu/cgi-bin/hgTables), AGO2 binding sites identified using iCLIP-Seq ^39^, PABPC1 and ELAVL1 bound RNAs identified using RIP-Seq (downloaded from https://www.ncbi.nlm.nih.gov/geo/; accession codes: GSM944519 and GSM944520), plus predicted miRNA target sites from TargetScan ^40^ for highly expressed miRNAs in LCLs according to miRNA measurements in Geuvadis ^2^, downloaded from http://www.ebi.ac.uk/arrayexpress/files/E-GEUV-3/GD452.MirnaQuantCount.1.2N.50FN.sample name.resk10.txt.

To compare the overlap of QTLs with the above-described genomic sites between groups of QTLs (for different QTL types and shared QTLs) we calculated for each group the enrichment as the ratio between the proportion of QTLs of that group overlapping a genomic site and the proportion of QTLs that overlap among QTLs not in that group (and not shared with them). The enrichment within genomic regions for QTLs in each cluster is calculated as above by comparing with QTLs in other clusters. For the overlap with different gene regions (Figures 2B and 4F), only overlaps of QTLs with the associated gene’s regions were considered.

### Trans-QTL associations for cis-QTLs of TFs, RBPs, and miRNAs

We determined nominal trans-associations using QTLtools ^37^ for cis-QTLs of transcription factors (TFs), RNA-binding proteins (RBPs) and miRNAs. We took TFs from ^41^ and RBPs from ^42^. We tested all cis-QTLs within the first conditional QTL signal of each regulatory factor for trans-associations with exon levels (trans-exQTLs), intron levels (trans-inQTLs), and exon-intron-ratios (trans-ex-inQTLs) of genes located on different chromosomes or at distances of more than five million nucleotides from the cis-QTL. Cis-QTLs for miRNA expression levels obtained in LCLs (Geuvadis data set without African, YRI, samples) were taken from ^2^. To account for multiple testing, we applied a nominal Bonferroni threshold ^43^. We considered a trans-association significant if the nominal p-value was smaller than 0.05 / ((number of tested TFs, RBPs or miRNAs) x (number of PCs explaining 95% of the variance in gene expression between individuals)). The number of PCs were 774 for exon levels, 795 for intron levels, and 806 for exon-intron ratios. In order to compare the proportions of different trans-QTL associations across different regulatory factors, we only considered trans-associations of genes for which all three trans-QTL types were testable. We discarded trans-associations for five genes that overlapped with pseudogenes of coding genes that were located within one million nucleotides of the tested QTLs. Although we did not count RNA-Seq reads overlapping more than one gene annotation, there could still be sequence similarities leading to mis-aligned reads between the pseudogene and the gene in the cis-window around the QTL.

### Colocalization of cis-QTLs with GWAS variants

We downloaded GWAS associations from different studies from the GWAS catalog (https://www.ebi.ac.uk/gwas/; version 1.0) ^44^. We considered GWAS variants with an association p-value < 1e^-8^, from studies carried out with European individuals or replicated with European individuals. This resulted in 53438 unique GWAS variants, of which 49711 variants were also genotyped in the CoLaus/Geuvadis data set. We performed the colocalization analysis between GWAS variants and top cis-QTLs using the regulatory trait concordance (RTC) method ^8^ implemented in QTLtools 1.3 ^37^, with 160 times sampling from simulated data. A colocalization was significant if the p-value of the RTC score was below 0.05.

### Clustering of colocalizing cis-QTLs based on relative effects

To make QTL effects comparable between QTL types and across variants, we first normalised the absolute effect sizes of each variant to its maximum among the three types of cis-effects. Then, we z-scored the normalised absolute effect sizes across all variants, separately for each of the three types of QTL effects. Hierarchical clustering of relative effect sizes was performed with scipy.cluster.hierarchy.linkage in Python ^45^ using the squared euclidean distance metrics and the weighted method for calculating distances between clusters.

## Availability of data and materials

exQTLs, inQTLs and ex-inQTLs (associated with exon and intron expression levels of genes, as well as their ratio, respectively) identified in this study are available as Supplementary Tables. The Geuvadis data set used in this study is publicly available, while the CoLaus data set is under restricted access and available upon request. All other data analysed in this study is publicly available from diverse sources, as indicated in the respective Methods sections.

## Competing interests

The authors declare no competing interests.

## Funding

This study was supported by the Swiss National Science Foundation (Grant No. FN 310030_152724/1).

## Acknowledgments

We thank Zoltan Kutalik for providing imputed genotypes of the CoLaus data set, and aSciStance Ltd for help in revising the manuscript.

## Authors’ Contributions

A.B. designed the project. A.B. carried out the computational analyses and prepared the results. A.B. and S.B. discussed the results. A.B. wrote the manuscript.

---

**Figure S1:**
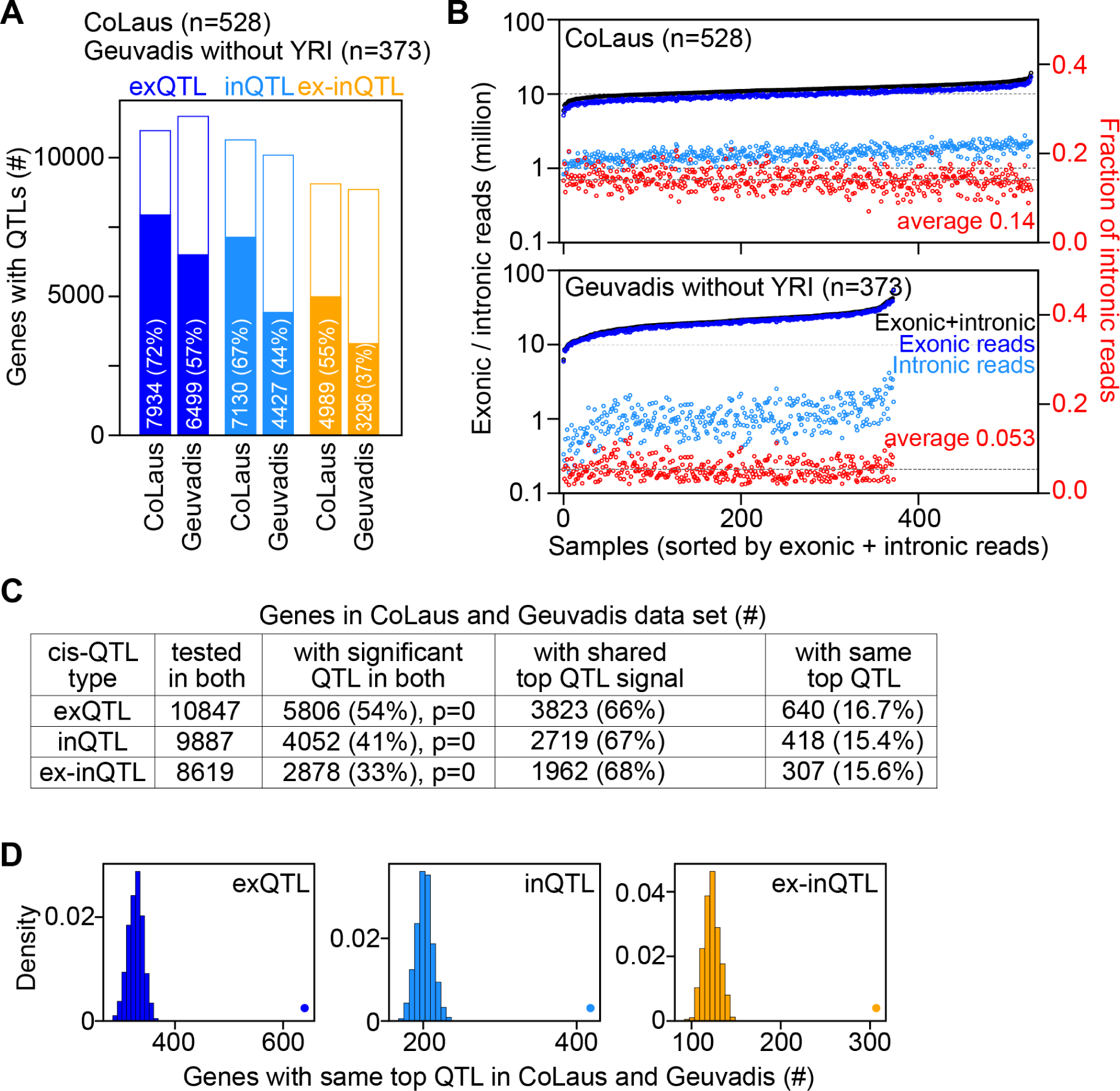
Replication of top cis-exQTLs, cis-inQTLs and cis-ex-inQTLs in the CoLaus and Geuvadis data sets. (A) Number and percentages of genes with cis-QTLs (filled bars) of tested genes (full bars) for exQTLs, inQTLs and ex-inQTLs for the CoLaus data set (n=528) and for Geuvadis data set, excluding samples from Afrikan individuals from Yoruba (YRI), (n=373). (B) Number of RNA-Seq reads mapping uniquely to exons, introns, or both for samples in the Colaus data set (top panel) and the Geuvadis, without YRI, data set (bottom panel), sorted by number of reads mapping to exons and introns. The second y-axis (right) indicates the fraction of reads mapping uniquely to introns (red circles). (C) Number of genes that were testable, had significant cis-QTLs, had shared cis-QTL signals, had identical top cis-QTLs in the CoLaus and Geuvadis without YRI data sets for each cis-QTL types. Percentages are relative to the number in the previous column. P values were calculated with a hypergeometric test. (D) The number of genes with identical top cis-QTLs in the CoLaus and Geuvadis without YRI data sets (indicated by a dot) is significantly larger, than expected when drawing randomly one cis-QTL from the QTLs sharing the top QTL signal (distribution for 100 random draws shown as histogram), for all types of cis-QTLs (p<1e-100; one sample T test compared with 100 random sets).

**Figure S2:**
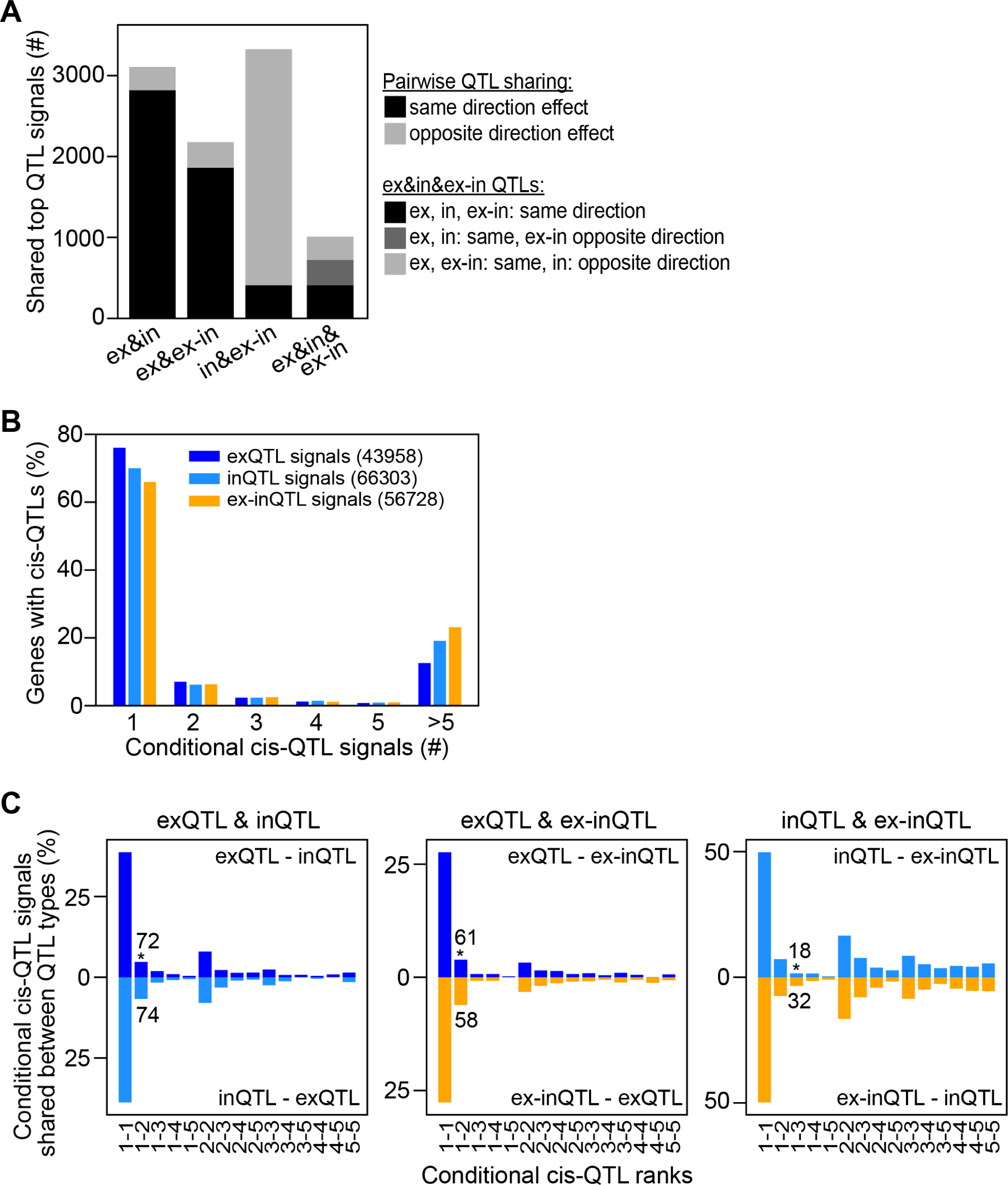
Sharing between top cis-QTLs and between conditional cis-QTLs. (A) Number of genes with shared top cis-QTL signals which have effects in the same or opposite direction. In case of sharing between all three QTL types, the numbers are further divided into all effects having the same direction, exQTL and inQTL effects having the same direction and the ex-inQTL effect the opposite direction, or exQTL and ex-inQTL effects having the same and the inQTL effect the opposite direction. (B) Percentage of genes with certain numbers of conditional cis-QTL signals (see Methods). The total number of conditional cis-QTL signals of each type is indicated in parenthesis. (C) Percentage of conditional cis-QTL signals of ranks 1 to 5 (indicated on the x-axis) that are shared between QTL types, as indicated on top of each panel. Bars in top direction show the comparison between ranks of two QTL types as indicated on top right, while bars in bottom direction show the comparison between the same QTL types, but with reversed ranks, indicated at bottom right. The number of shared conditional QTL signals is indicated in case of significantly different proportions. *, p<0.05, Fisher’s exact test.

**Figure S3:**
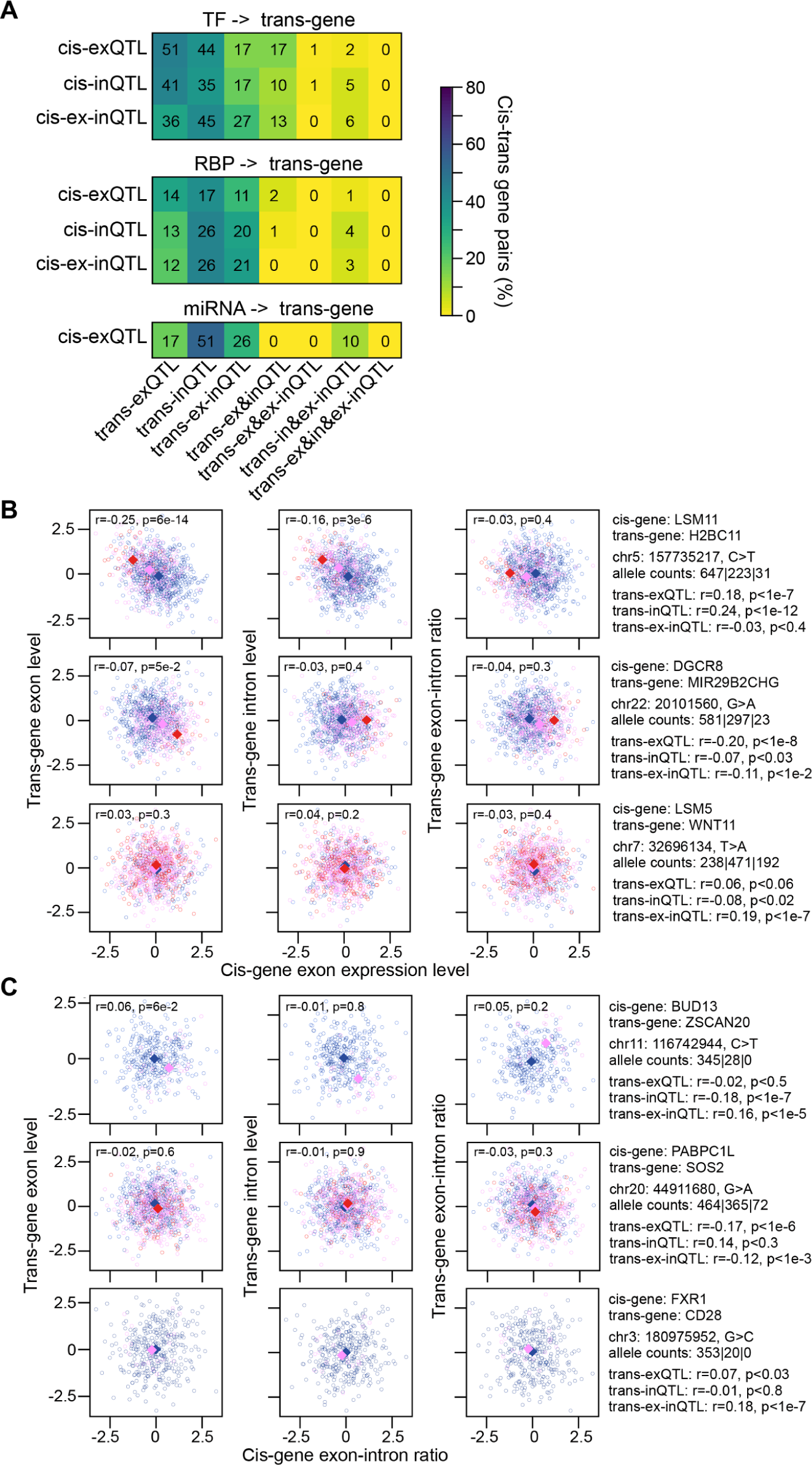
Trans-QTL associations for cis-QTLs of transcription factors, RNA-binding proteins, and miRNAs. (A) Percentage of unique cis-trans-QTL gene pairs (indicated on the y axis) for cis-exQTLs, cis-inQTLs and cis-ex-inQTLs of transcription factors (TFs; top panel) and RNA-binding proteins (RBPs; middle panel) and miRNAs (bottom panel). The number of unique cis-trans-QTL gene pairs of each type are indicated. (B) Examples for cis-regulated RBPs (one per row) associated with another gene in trans through cis-exQTLs. Shown are scatter plots of the RBP’s normalised exon levels (x-axis) and the normalised exon levels (left panels), intron levels (middle panels) and their ratio (right panels) of the trans-associated gene (y-axis). Information on the involved genes, QTL variants and trans-QTL associations are indicated on the right. Correlation coefficients between the expression levels/ratios of the RBPs and their trans-associated genes are indicated inside each panel. Circles of different colors represent individuals with different genotypes, and colored squares indicate the median values for individuals with that genotype. (C) Same as (B), but for associations in trans through cis-ex-inQTLs, and normalized exon-intron-ratios on the x-axis.

**Figure S4:**
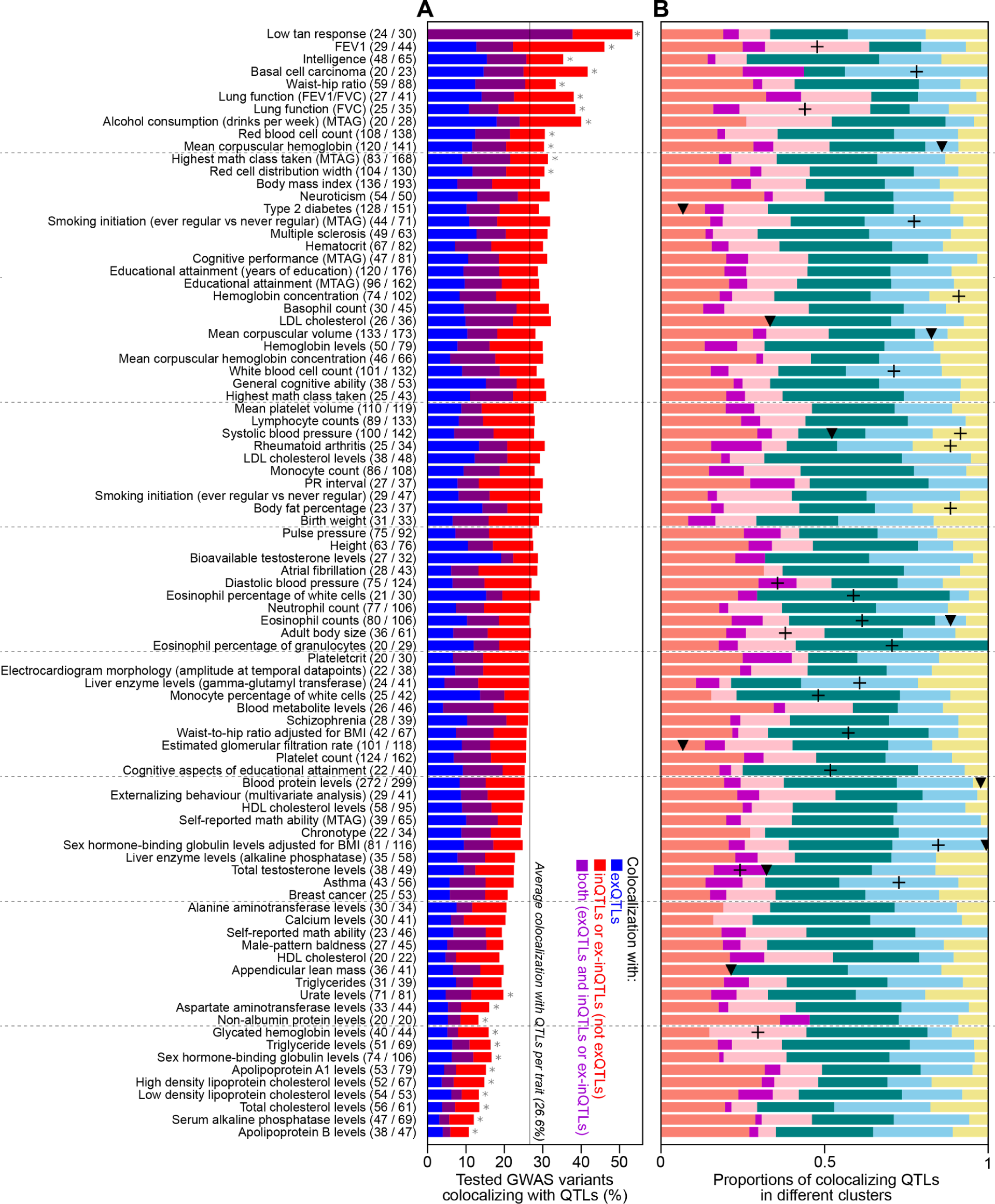
Colocalization between top cis-QTLs and GWAS variants for 89 traits with at least 20 colocalizations. (A) Proportion of tested GWAS variants colocalizing with top cis-QTLs for 89 GWAS traits (indicated at right) with at least 20 colocalizations. GWAS traits are sorted by the significance of the colocalization with cis-QTLs (calculated using Fisher’s exact test compared to all 89 GWAS traits, *: p<0.05). The number of colocalizing GWAS variants and the number of colocalizing cis-QTLs are indicated in parenthesis. The percentage of GWAS variants colocalizing with only exQTLs, not with exQTLs, and with exQTLs and another QTL type are indicated in blue, red, and purple, respectively. (B) Proportion of colocalizing top cis-QTLs with quantified effects on all three gene expression measures that are assigned to certain clusters according to their relative effect sizes (shown in Figure 4D). A plus or triangle symbol indicates an enrichment or depletion, respectively, in cis-QTLs of a certain cluster for a given trait compared to colocalizing cis-QTLs of all traits assigned to clusters (p<0.05, calculated using Fisher exact test).

**Figure S5:**
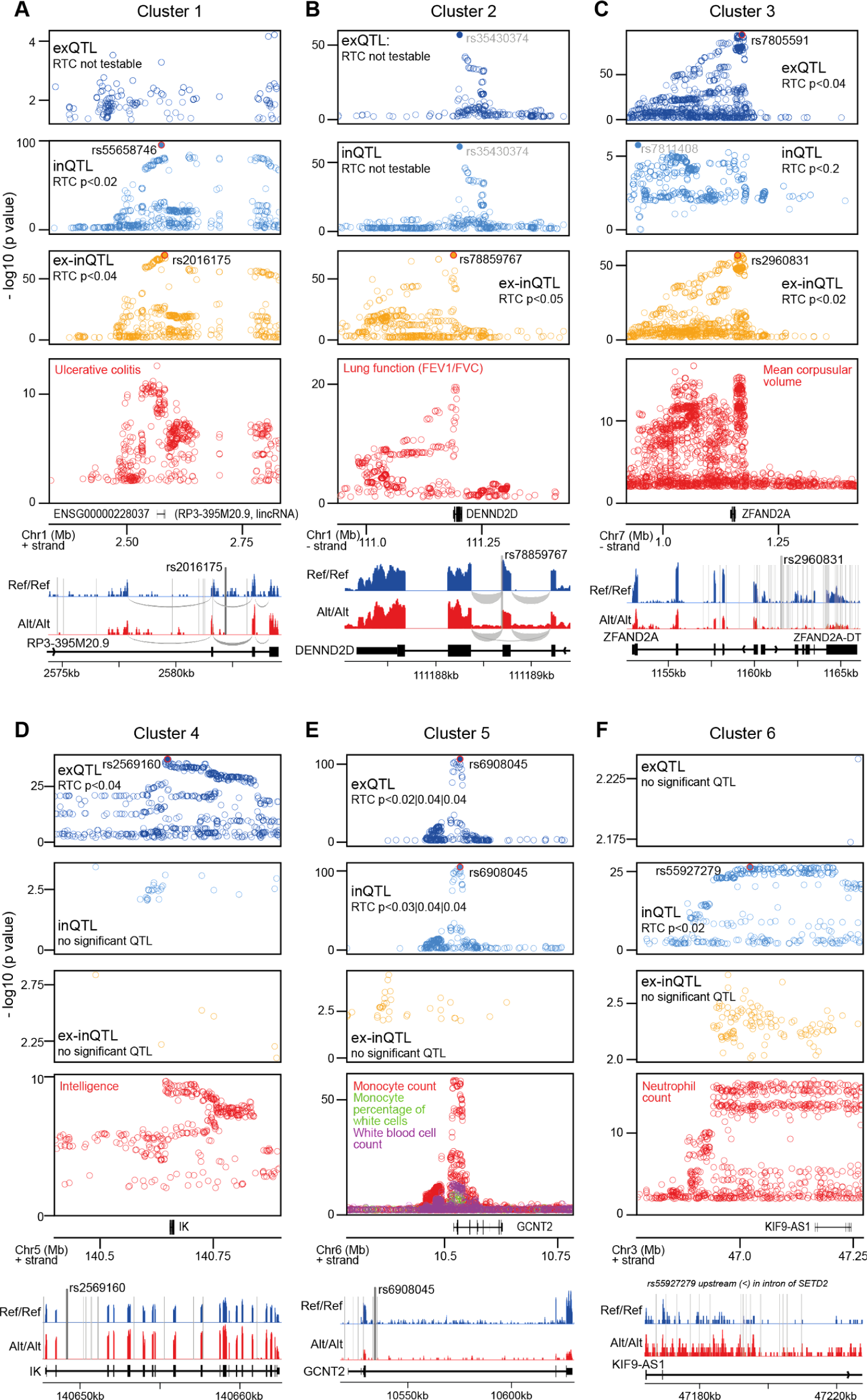
Further examples for cis-QTLs from different clusters colocalizing with GWAS variants. In each subfigure, the top three panels show, in a region of 500kb around the top-cis-QTL (x-axis), the -log10 nominal p-values (<0.01) for cis-QTL associations with exon levels (dark blue), intron levels (blue) and exon-intron-ratios (orange), and the fourth panel shows the -log10 p-values for the GWAS trait associations (red). The rsID of the top cis-QTL(s) are indicated (in black, if colocalized with the GWAS trait variants, or in grey if not). Colocalization via the RTC method (see Methods) is only testable for QTLs and GWAS variants within the same recombination hotspot. The bottom panel shows examples for RNA-Seq read distributions at the associated gene (or a region thereof) from two homozygous individuals, one with reference (Ref/Ref; blue) and one with alternative (Alt/Alt; red) genotype, for the cis-QTL variant. The positions of the top cis-QTL as well as cis-QTLs sharing the QTL signal are indicated with thick or thin lines, respectively. (A) A cis-QTL from cluster 1, associated with the long, intergenic non-coding RNA (lincRNA) RP3-395M20.9 and colocalizing with a GWAS variant for ulcerative colitis. The gene has been connected to autoimmune diseases before ^46^. The top cis-QTLs for intron levels and exon-intron-ratios are shared. The top cis-ex-inQTL is located in the second last intron of the lincRNA and appears to increase the rate of splicing of that intron (bottom panel). (B) A cis-QTL from cluster 2, associated with the exon-intron-ratio of DENND2D, a guanyl-nucleotide exchange factor promoting the exchange of GDP to GTP, and colocalizes with GWAS variants for the lung function trait FEV1/FVC. The expression of this gene is also affected by a shared QTL affecting exon and intron levels, which did not colocalize with variants for this GWAS trait. The top cis-ex-inQTL is at the 5’ end of the second last intron, between two coding exons, and appears to slow down splicing of the upstream exon, with a probability for skipping that entire coding exon, which is almost 90 nucleotides long. (C) A cis-QTL from cluster 3, associated with ZFAND2A and colocalizing with a GWAS variant for mean corpuscular volume. The top cis-QTLs for exon levels and exon-intron-ratios are shared, while the cis-QTL signal for intron levels is different and does not colocalize with the GWAS trait variants. The top cis-ex-inQTL is located in the first intron of the upstream non-coding gene, ZFAND2A-DT, which is annotated as an divergent transcript of ZFAND2A. The top cis-ex-inQTL shows a reduction in expression of both genes. It can not be excluded that these genes, that appear to be co-transcriptionally regulated with histone acetylation marks around their gene starts, have additional RNA-interactions after transcription. (D) A cis-QTL from cluster 4, associated with IK and colocalizing with a GWAS variant for intelligence. The top cis-exQTL is located in the second intron of the gene, and likely increases the transcription of that gene. (E) A cis-QTL from cluster 5, associated with GCNT2 and colocalizing with GWAS variants for three related traits, monocyte count, monocyte percentage of white cells, and white blood cell count. The top cis-QTL is identical for exon and intron levels. It is located in the third (longest) intron and likely reduces transcription of that gene, detectable at exon and intron levels. (F) A cis-QTL from cluster 6, associated with KIF9-AS1 and colocalizing with a GWAS variant for neutrophil count. There are no significant cis-QTLs for exon levels and exon-intron-ratios. The colocalizing top cis-QTL for intron levels is located in the intron of an upstream gene, SETD2, and likely increases transcription slightly, without an increase in splicing, such that it is only detectable at intron levels.

## Supplementary Text

### Replication of top cis-QTLs in CoLaus and Geuvadis data sets

To evaluate the reproducibility of cis-QTLs for exon and intron levels and for their ratio, we mapped such cis-QTLs in two independent data sets, both consisting of RNA-Seq data from lymphoblastoid cell lines (LCLs) and genotypes from European individuals. In particular, we analysed 528 samples of CoLaus data set ^23^, and 373 samples of the Geuvadis data set ^2^, excluding samples from African individuals from Yoruba (YRI). We separately counted RNA-Seq reads mapping uniquely to either exons or introns, and quantified exonic and intronic expression levels for each gene, as well as the ratio of these (see Methods for details). Genes were tested for QTL associations if the median number of uniquely mapping RNA-Seq reads was at least 10 across individuals in a data set. We detected significant (FDR < 5%) exQTLs, inQTLs and ex-inQTLs in cis for 72%, 67% and 55% of testable genes, respectively, in the Colaus data set, and for 57%, 44% and 37% of testable genes, respectively, in the Geuvadis (without YRI) data set (Figure S1A).

Overall, the percentage of genes with QTLs was lower for the Geuvadis (without YRI) data set than for the CoLaus data set, likely due to the smaller sample size (71% of CoLaus data set). For both data sets, the percentage of genes with inQTLs was lower compared to the percentage of genes with exQTLs, likely because of the lower number of intronic RNA-Seq reads compared to exonic RNA-Seq reads, and thus a lower accuracy in the measured expression levels of introns compared to exons (Figure S1B). The percentage of genes with inQTLs and ex-inQTLs was particularly lower in the Geuvadis (without YRI) data set, likely due to a lower proportion of RNA-Seq reads mapping to introns (on average 0.14 for CoLaus versus 0.053 for Geuvadis data set; Figure S1B).

Of the genes that were tested for cis-QTL associations in both data sets, 54%, 41% and 33% had significant exQTLs, inQTLs, and ex-inQTLs in both data sets, which was a significant overlap (p=0, hypergeometric test; Figure S1C). Of these, the top cis-QTLs (most significantly associated genetic variant) for a gene were shared, i.e. within the leading QTL signal (all genetic variants with similar, but slightly less significant, association as the top QTL) and with consistent effect directions, between both data sets, for 66% of exQTLs, 67% of inQTLs, and 68% of ex-inQTLs (see Methods for details on cis-QTL sharing). The percentage of shared cis-QTLs was similar for different types of QTLs, indicating similar reproducibility for all types of QTLs. The fraction of shared cis-QTLs with identical positions in both data sets was 16.7%, 15.4% and 15.6%, which was significantly larger than expected, when randomly picking one variant among the variants sharing the leading QTL signal (Figure S1D).

In summary, despite differences in sample size and in the fraction of intronic reads, all types of QTLs are equally reproducible across the two independent CoLaus and Geuvadis (without YRI) data sets.

## Notes

### Competing Interest Statement

The authors have declared no competing interest.

